# A circadian behavioral analysis suite for real-time classification of daily rhythms in complex behaviors

**DOI:** 10.1101/2024.02.23.581778

**Authors:** Logan J. Perry, Blanca E. Perez, Larissa Rays Wahba, KL Nikhil, William C. Lenzen, Jeff R. Jones

**Affiliations:** Department of Biology, Texas A&M University, College Station, TX; Department of Biology, Washington University in St. Louis, St. Louis, MO; Institute for Neuroscience, Texas A&M University, College Station, TX; Center for Biological Clocks Research, Texas A&M University, College Station, TX

## Abstract

Measuring animal behavior over long timescales has been traditionally limited to behaviors that are easily measurable with real-time sensors. More complex behaviors have been measured over time, but these approaches are considerably more challenging due to the intensive manual effort required for scoring behaviors. Recent advances in machine learning have introduced automated behavior analysis methods, but these often overlook long-term behavioral patterns and struggle with classification in varying environmental conditions. To address this, we developed a pipeline that enables continuous, parallel recording and acquisition of animal behavior for an indefinite duration. As part of this pipeline, we applied a recent breakthrough self-supervised computer vision model to reduce training bias and overfitting and to ensure classification robustness. Our system automatically classifies animal behaviors with a performance approaching that of expert-level human labelers. Critically, classification occurs continuously, across multiple animals, and in real time. As a proof-of-concept, we used our system to record behavior from 97 mice over two weeks to test the hypothesis that sex and estrogen influence circadian rhythms in nine distinct home cage behaviors. We discovered novel sex- and estrogen-dependent differences in circadian properties of several behaviors including digging and nesting rhythms. We present a generalized version of our pipeline and novel classification model, the “circadian behavioral analysis suite,” (CBAS) as a user-friendly, open-source software package that allows researchers to automatically acquire and analyze behavioral rhythms with a throughput that rivals sensor-based methods, allowing for the temporal and circadian analysis of behaviors that were previously difficult or impossible to observe.

## Introduction

Understanding the genetic, neural, and ethological mechanisms that temporally organize behavior is a fundamental goal of fields including circadian biology, neuroscience, and ecology. However, the temporal analysis of behavior has been largely limited to behaviors that can be accurately measured with low latency and at high throughput. For instance, optical and electrical sensors enable such analysis of eating, drinking, and locomotor behaviors. These behaviors have been widely studied (although usually independently) at high temporal resolution for experimental durations of weeks, months, or even years (Schwartz and Zimmerman, 1990; Jud et al., 2005; Pendergast et al., 2013; Yamanaka et al., 2013; Metzger et al., 2020). Other more complex behaviors such as rearing, nesting, and grooming can often be measured simultaneously using video recording and manual behavior scoring by trained human observers (van der Veen et al., 2008; Gaskill et al., 2009; van Oosterhout et al., 2012; Fujita et al., 2017; Robinson-Junker et al., 2018; Shuboni-Mulligan et al., 2021). Over long timescales, however, this method becomes impractical because video acquisition inevitably outpaces human labeling, leading to an ever-increasing latency between data acquisition and data analysis. For example, it may take an observer less than a minute to label behaviors in a minute long video recording but, due to the tedium of the task and the limits of the human attention span, it would take that observer much longer than a week to classify behaviors in a week-long video recording (Segalin et al., 2020; Muller et al., 2021). Consequently, these behaviors have been studied infrequently at low temporal resolution for limited experimental durations, such as hourly over the course of a single day. To better understand how animal behavior changes over time, ethologically relevant behaviors (regardless of “measurability”) must be measured in individual animals simultaneously, automatically, and, critically, in real time over multiple days and conditions at high temporal resolution (Peters et al., 2015; Grieco et al., 2021; Kahnau et al., 2023).

Over the last two decades, methods have been developed to classify animal behavior from video recordings, ranging from computer vision algorithms, such as centroid tracking, to machine learning-based approaches that use markerless pose estimation (e.g., DeepLabCut or DLC) or raw pixel values (e.g., DeepEthogram or DEG) to quantify behavior (Mathis et al., 2018; Pereira et al., 2020; Zhang et al., 2020; Bohnslav et al., 2021). While previous studies have used machine learning to analyze temporal variation in more naturalistic “home cage” behaviors, these methods have faced several challenges (Steele et al., 2007; Goulding et al., 2008; Jhuang et al., 2010; Adamah-Biassi et al., 2014; Salem et al., 2015). For instance, existing methods tend to disregard long-range temporal information by simplifying analysis to frame-wise positional and motion values. A more holistic approach is needed to capture the temporal dynamics of both the recorded animal and potentially changing in-scene objects on both short and long time scales (Xie et al., 2017). Additionally, existing methods are often constrained by specific environmental conditions, such as video perspective, lighting condition, or subject coloration, which greatly limits their applicability. The development of an adaptable, condition-agnostic system is therefore essential for robust temporal analysis. Perhaps most importantly, existing methods have not been typically used for real-time behavior analysis and have not been used to analyze behavior on a “circadian” timescale of days to weeks or longer. Together, these challenges have prevented the widespread adoption of machine learning classification to the long-term or circadian analysis of behavior. This is likely exacerbated by the lack of user-friendly tools that facilitate acquisition, training, validation, and analysis of key behavioral metrics. To solve this problem, we developed a streamlined approach that allows users to extend modern deep learning methods to emulate the functionality of traditional sensor-based analyses of behavior.

Here, we introduce a versatile, high-throughput, real-time behavior acquisition and analysis pipeline for the temporal analysis of behavior. To do this, we created the software infrastructure to automatically acquire behavior data from video recording streams in real time, in parallel (here, from 24 mice simultaneously) for an essentially unlimited experimental duration (here, for up to two weeks continuously at 10 frames per second). Next, we developed a joint long short-term memory and linear layer model to integrate the visual and motion features output by DINOv2, a state-of-the-art self-supervised computer vision feature extractor. Finally, we combined this model with our recording pipeline to facilitate indefinite recording and behavioral analysis. As a proof-of-concept, we trained our classification model to identify nine home cage behaviors (eating, drinking, rearing, climbing, digging, nesting, resting, grooming, and locomotion; (Garner, 2017)) in male and female mice to test the hypothesis that sex and estrogen influence circadian rhythms in home cage behaviors. Previous studies have identified subtle sex differences in wheel-running activity rhythms (Krizo and Mintz, 2014; Joye and Evans, 2022). However, despite the global regulation of behavior by the circadian system, sex differences in other behavioral rhythms have not yet been identified due to technological limitations. Our automatic inference system allowed us to discover novel sex- and estrogen-dependent differences in the phase and amplitude of several behaviors, including, notably, digging and nesting rhythms. Finally, we developed our DINOv2 model and automatic inference software into a user-friendly, open-source Python package called CBAS, the “circadian behavioral analysis suite.” CBAS allows researchers to automatically acquire and analyze behavioral rhythms with a throughput that greatly exceeds manual video labeling and rivals sensor-based methods.

## Results

### Machine learning classification of behaviors approaches human level performance

We first recorded continuous videos of individually-housed mice for >24 h at 10 fps in both a 12 h:12 h light:dark (LD, where dark is defined as dim 850 nm infrared light) cycle and in constant darkness (DD) (**Fig. 1a**). From these videos, we used strict criteria (**Supplementary Table 1**) to define and manually label nine ethologically-relevant behaviors that encompass the majority of an individual singly-housed mouse’s daily behavioral repertoire, including maintenance, exploratory, and inactive behaviors: eating, drinking, rearing, climbing, grooming, exploring, digging, nesting, and resting (**Fig. 1b**) (Garner, 2017). For each behavior, we identified the average length of time for a “bout,” or behavioral instance (**Fig. 1c**). This allowed us to generate balanced training and test sets from segments of videos sampled from 30 mice that contained a balanced number of unique instances of each behavior (**Fig. 1d**). To control for lighting conditions, we sampled video segments such that there was a balanced representation of each behavior during both the animal’s active and inactive phases.

**Figure 1.**
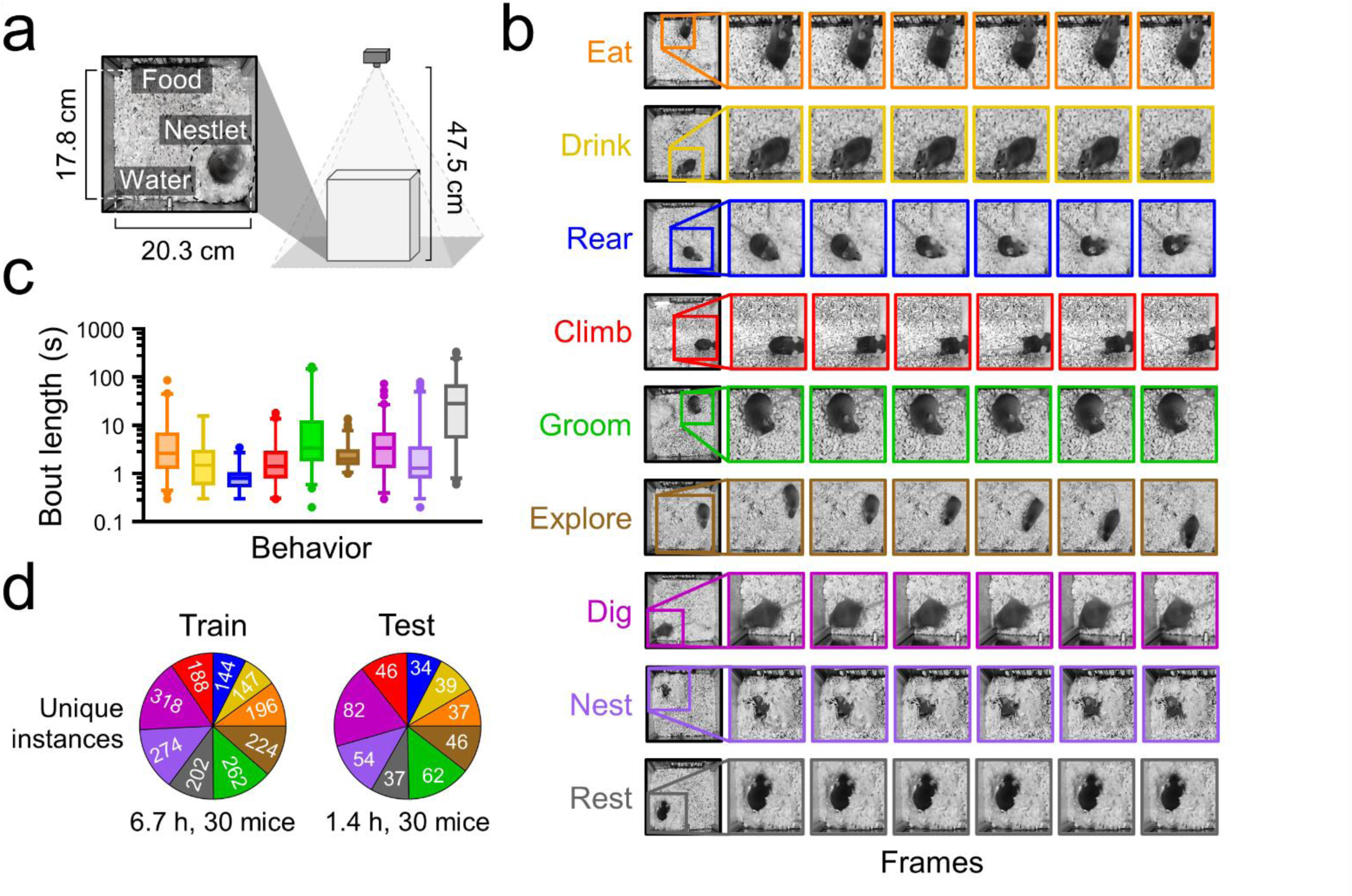
Recording and classification standardization of nine home cage behaviors. **a)** Schematic of the home cage recording setup. **b)** Representative examples of individual frames depicting each of nine behaviors (eating, orange; drinking, yellow; rearing, blue; climbing, red; grooming, green; exploring, brown; digging, magenta; nesting, purple; resting, gray). First frame depicts the behavior occurring in the full field of view, subsequent frames are zoomed in to better illustrate behaviors. **c)** Bout length (duration of a behavioral instance) for each behavior within a maximum window size of 360 s. n ≥ 38 bouts from 29 to 30 mice per behavior. Box and whiskers depict median and interquartile range. **d)** Number of unique instances of each behavior in the 8.1 h human-labeled dataset broken down by training and test sets.

We next used these training and test sets to train a previously published deep learning behavior classification model, DeepEthogram (DEG) (Bohnslav et al., 2021), and our own DINOv2+ model (**Fig. 2a**). We constructed DINOv2+ using the state-of-the-art DINOv2 vision transformer model (Oquab et al., 2023) as a “frozen,” or immutable, feature extractor ‘backbone’ with a trainable joint long short-term memory and linear layer classification network ‘head.’ DEG and DINOv2+ are each capable of producing behavior classifications from a video frame’s raw pixel values as binary output matrices (“ethograms”) that indicate if a behavior is present or absent in a given frame. This temporally sequenced ethogram output is ideal for quantifying behavioral rhythms because it is readily analyzed using field-standard circadian analysis methods that are optimized for time series data. However, while DEG is trained using a supervised learning scheme, the backbone feature extractor of our DINOv2+ model is pretrained using a self-supervised approach that has been shown to be more generalizable (Tendle and Hasan, 2021). Thus, training and testing both models allows us to directly compare the performance of these two different underlying learning schemes on visual feature extraction.

**Figure 2.**
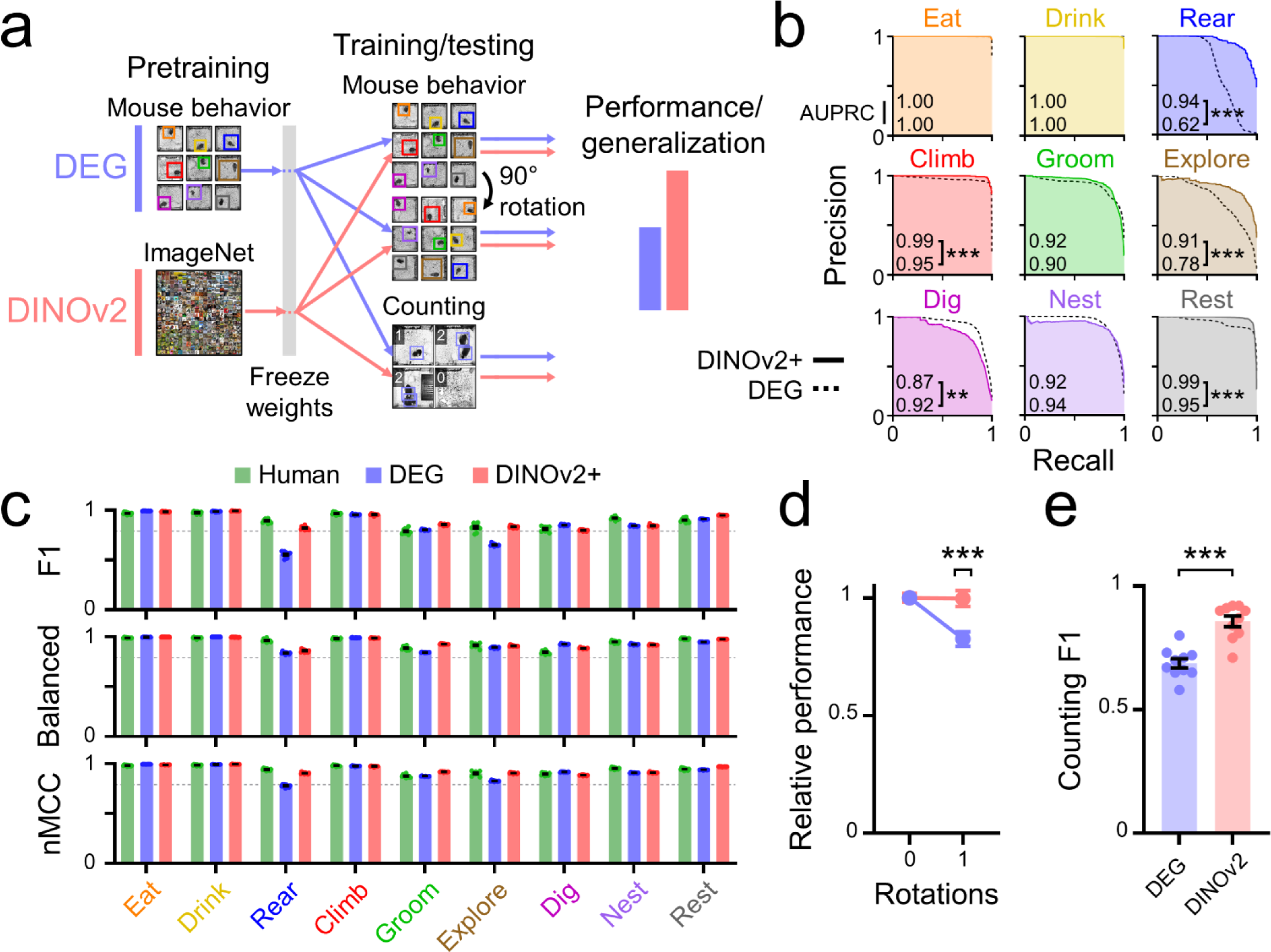
DINOv2+ approaches expert-level performance on behavior classification. **a)** Schematic of performance and generalization tests. Features from a frozen pretrained DeepEthogram (DEG) model and a frozen pretrained DINOv2 model were used to evaluate the ability of each visual feature extractor to successfully classify mouse behavior using our DINOv2+ joint LSTM and linear layer model head (performance; **Figs. 2b,c**), classify mouse behavior on behavior frames rotated 90° using a single layer linear network head (generalization; Fig. 2d), and count the number of mice in a cage using a single layer linear network head (generalization; Fig. 2e). **b)** Precision-recall curves for each behavior calculated for the DINOv2+ (colored lines) and DEG (dashed lines) models by varying the decision threshold of each binary classifier. Shading depicts the area under the precision-recall curve (AUPRC) for each behavior for each model. Bootstrap test; **, p < 0.01; ***, p < 0.001. **c)** Performance metrics for each behavior calculated for a trained human classifier (green), the DEG model (blue), and the DINOv2+ model (red). n = 10 sets of 1,000 randomly sampled test set frames per behavior. Dashed line depicts a predefined performance threshold of 0.80. Lines and error bars depict mean ± SEM. F1, F1 score; nMCC, normalized Matthews correlation coefficient. **d)** Relative performance for the DEG (blue) and DINOv2 (red) pretrained models when tested on a rotated version of a baseline behavior sequence test set using a single layer linear network head on top of the baseline models. **e)** F1 score calculated for both DEG and DINOv2 on a classification task involving counting the number of mice in a cage using a single layer linear network head on top of the baseline models.

If we wanted to use our models to automatically infer days of video – millions or potentially billions of frames that would never be seen by a human – it was critical that our models were extensively validated. Model performances quantified across all measured behaviors of existing commercial (e.g., HomeCageScan) and non-commercial methods used for the temporal analysis of home cage behaviors are either unreported or, typically, mediocre. Thus, after training our models, we performed rigorous validation of the model’s predictions on our labeled test sets with stringent model performance thresholds. Importantly, we did not adjust model hyperparameters based on our model’s test set performance. We compared the performance of our DEG and DINOv2+ models with that of a trained human classifier. Each of these groups were given a 15-31 frame (1.5-3.1 s) window to predict behaviors from our test set.

First, because most machine learning performance metrics require us to define a specific threshold value at which behavior probabilities are converted into a binary prediction, we generated precision-recall curves across different probability thresholds for our DEG and DINOv2+ models (**Fig. 2b**). We did not generate human classifier precision-recall curves because in our training set human labels are inherently binary, not probabilistic. We found that the areas under the precision-recall curves (AUPRC, a summarization of model performance as a function of probability cut-off threshold) for our DINOv2+ model greatly outperformed DEG on rearing and exploring behaviors and slightly, but significantly, outperformed DEG on climbing and resting behaviors. DEG slightly, but significantly, outperformed DINOv2+ on digging behavior classification.

Next, we used multiple demanding metrics (F1 score, balanced accuracy, and normalized Matthews correlation coefficient; (Brzezinski et al., 2020; Chicco and Jurman, 2020; Grandini et al., 2020)) to test the performance of each of our models (**Fig. 2c**). Our predetermined criteria for a “successful” model was a score of at least 0.80 for each behavior on each metric. A successful model would also ideally meet or exceed the performance of a trained human classifier labeling the same test set. Our DINOv2+ model met or exceeded our predefined threshold on all performance metrics, whereas our DEG model failed to meet this F1 and nMCC score threshold for rearing and exploring behaviors. Notably, DINOv2+ exhibited greater performance than DEG even on behaviors that had F1, balanced accuracy, or nMCC scores of >0.80, such as grooming and resting. DINOv2+ also met or exceeded human classifier performance on metrics for most behaviors, including eating, drinking, climbing, grooming, exploring, digging, and resting, while DEG only met or exceeded human classifier performance on eating, drinking, climbing, and digging behaviors. Together, these results demonstrate that our DINOv2+ model’s performance on our test set approaches that of expert-level human classifiers. These performance results confidently indicate a high level of reliability of our model, which would allow us to perform behavior inference on a circadian timescale of days to weeks.

Finally, to assess the differences between supervised and self-supervised learning approaches in DEG and DINOv2, we trained two additional linear probe heads on top of the frozen outputs of a pretrained DEG model and the DINOv2 model. First, we trained a linear probe to classify our nine mouse behaviors using a training and test set comprising mouse behavior frames that were simply rotated 90 degrees from the original orientation of each frame used to pretrain the DEG model (**Fig. 2d**). We found that rotation had a negligible impact on the DINOv2 model’s performance. However, surprisingly, our DEG model’s performance dropped nearly 20% after a single rotation, even though DEG uses rotation as part of its image augmentation process (Bohnslav et al., 2021). Next, we trained a linear probe on a completely novel task in which both models must count the number of mice in a given frame using a training and test set comprising video frames containing zero, one, or two mice in their home cage with and without the presence of a running wheel (**Fig. 2e**). Unsurprisingly, because DINOv2 is a foundational model that can be adapted to a wide range of classification tasks, DINOv2 greatly outperformed DEG on this counting task. These results demonstrate the difference in visual feature robustness between supervised and self-supervised learning schemes and strongly suggest that the DINOv2 model can serve as a powerful pretrained backbone for a wide variety of classification tasks.

### Behavior classification occurs in real time

Regardless of our DINOv2+ model’s exceptional performance, the complexity of machine learning models often translates into poor usage speeds and low throughput in practice. Using DINOv2+ as an enhancement to (or replacement for) traditional sensor-based behavior analysis requires us to use it to infer videos in real time. That is, a video clip of n second duration must be recorded, processed, and automatically inferred by DINOv2+ before n seconds have elapsed and the next video segment is ready to be processed and inferred. To match the high throughput of sensor-based analysis (e.g., many running wheels can be recorded in parallel), we also need to be able to record, process, and automatically infer behaviors from videos recorded from multiple mice simultaneously. To solve this problem, we developed a hardware and software pipeline that allowed us to automatically and continuously record and infer behaviors in real time from up to 24 mice in parallel (**Fig. 3a**). Our system comprises power over ethernet (PoE) IP cameras connected in parallel to Gigabit switches. These switches stream video data that is binned into constant length time segments onto dedicated machine learning computers for inference and network-attached storage devices for backup.

**Figure 3.**
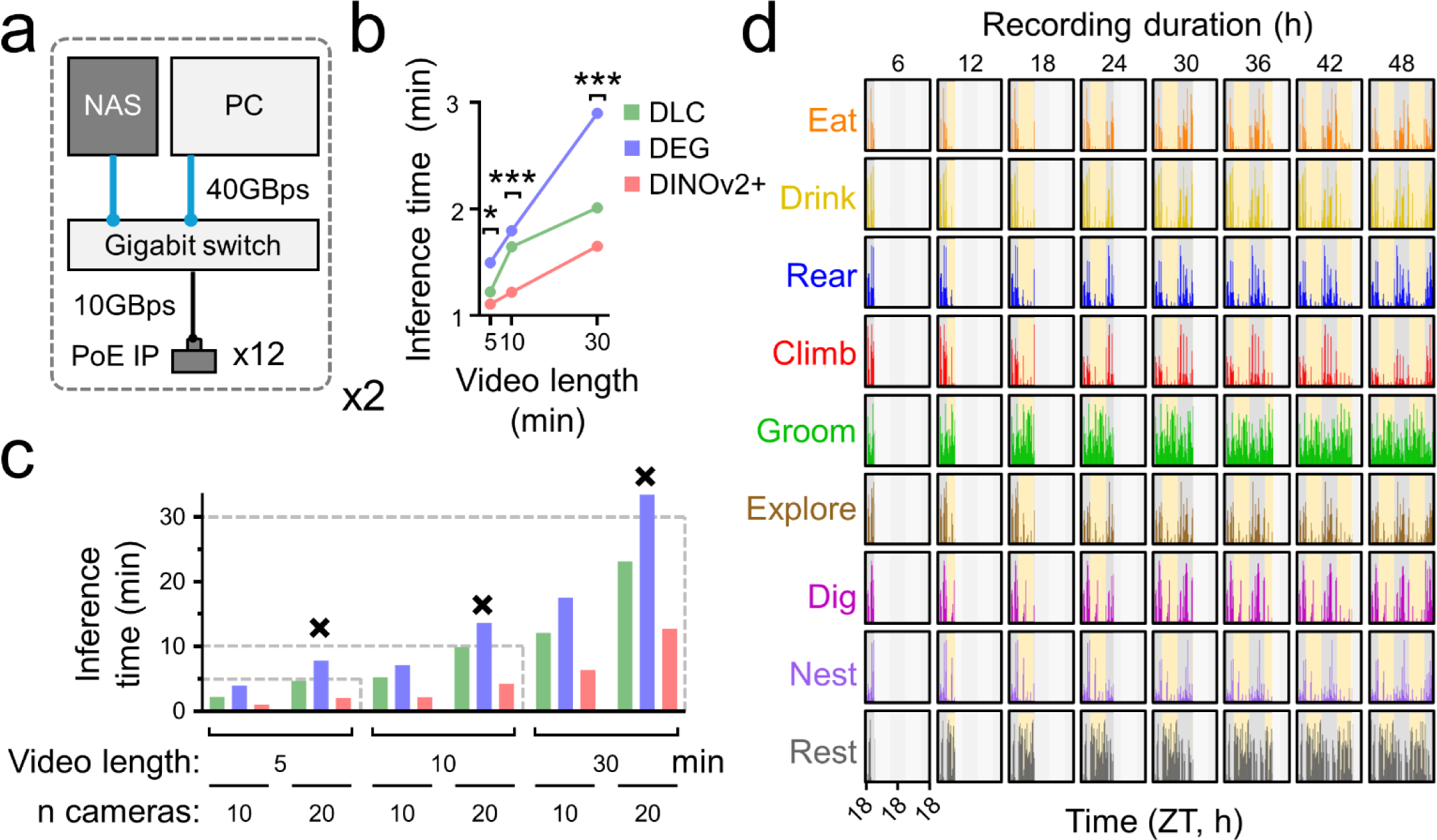
DINOv2+ allows for real-time behavior classification. **a)** Schematic of the real-time video recording, processing, and inferring system comprising two sets of 12 PoE (power over ethernet) IP cameras networked to a switch that passes streaming video data to a machine learning computer for video inference and a network-attached storage device for video backup. **b)** Single-video inference times for video segments of various lengths calculated for a skeletal pose estimation model without behavior classification (green, DLC), DEG (blue), and DINOv2+ (red). n = 3 replicates per model. Two-way ANOVA with post-hoc Tukey’s multiple comparison’s test; *, p < 0.05; ***, p < 0.001. **c)** Inference times for combinations of video segment length and number of cameras used to simultaneously stream video segments calculated for each model. Dashed lines depict the times at which inference time equals the length of the video segment. Failure of real-time inference for a particular combination of segment length, camera number, and inference model is represented by a black X above the bar. **d)** Representative activity profiles for each behavior from an individual mouse recorded in a 12 h:12 h light:dark (LD) cycle for 48 h. 30 min segments of continuously recorded video were automatically processed, inferred, and plotted over the duration of the recording, “filling in” over time. For visualization, plots shown here are only updated every 6 h. ZT, zeitgeber time.

Before using our system, we needed to identify the video segment length (in minutes) such that video data from x cameras can be inferred within that temporal window. To do this, we first calculated the single camera inference times for several potential models including DEG, DINOv2+, and, for comparison, a skeletal pose estimation model without behavior classification (DLC) (**Fig. 3b**) (Mathis et al., 2018). We found that while all models were able to infer video data from a single camera within each of the temporal windows tested, DINOv2+ was significantly faster at video inference than either DEG or DLC.

Next, to apply real-time inference to multiple animals in parallel, we first needed to identify the maximum number of cameras that could infer behaviors simultaneously within a reasonable video segment length. We again used DEG, DINOv2+, and DLC to calculate behavior inference times for various combinations of time segment lengths (5 min, 10 min, or 30 min) and numbers of cameras used to simultaneously stream video segments (10 or 20; **Fig. 3c**). We found that all models were able to infer video data from 10 cameras simultaneously regardless of video segment length. However, when our DEG model was used to infer video data from 20 cameras simultaneously, inference time exceeded the length of the video segment regardless of video segment length. This indicated a failure of real-time inference. In addition, our DINOv2+ model was significantly faster at video inference than either DEG or DLC at all time segment length and camera number combinations tested. Based on these results (and our experimental setup in which our behavior cabinets can hold a maximum of 12 mouse cages each), we chose to proceed with using our DINOv2+ model to infer videos with a video segment length of 30 min on a system comprising two sets of 12 cameras networked to individual machine learning computers (**Fig. 3a**). To test the efficacy of our system, we recorded videos from 24 mice simultaneously over 48 h in a 12h:12h LD cycle (**Fig. 3e, Supplementary Video 1**). Our system was able to successfully process video clips, infer behaviors, and plot time series activity profiles for each behavior over the duration of the recordings, “filling in” over time similarly to how wheel-running activity profiles are plotted by commercial circadian activity monitoring software. Together, these results demonstrate that our model can be used to automatically and continuously classify behaviors from multiple animals for an essentially unlimited experimental duration.

### Sex influences circadian rhythms in home cage behaviors

Next, we applied our DINOv2+ model and automatic inference system to a fundamental question in circadian biology: how do sex and estrogen influence circadian rhythms in behavior? Subtle sex differences in wheel-running activity rhythms have been previously identified (Lee et al., 2004; Kuljis et al., 2013, 2016; Krizo and Mintz, 2014; Joye and Evans, 2022; Anderson et al., 2023). However, because of technological constraints, whether (and how) males and females differ in other behavioral rhythms is unknown. To address this problem, we continuously recorded videos, inferred behaviors, and generated actograms (a field-standard method of plotting activity profiles over multiple days) from male (n = 24) and female (n = 27) mice over 5 d in LD and over 5 to 9 d in DD (**Figs. 4a, b**). Female mice underwent estrous staging prior to beginning recording, allowing us to sort them into groups adjusted such that their actograms were aligned by their first day of proestrus. We used these actograms to determine key circadian properties of each behavioral rhythm including phase, amplitude, and period.

**Figure 4.**
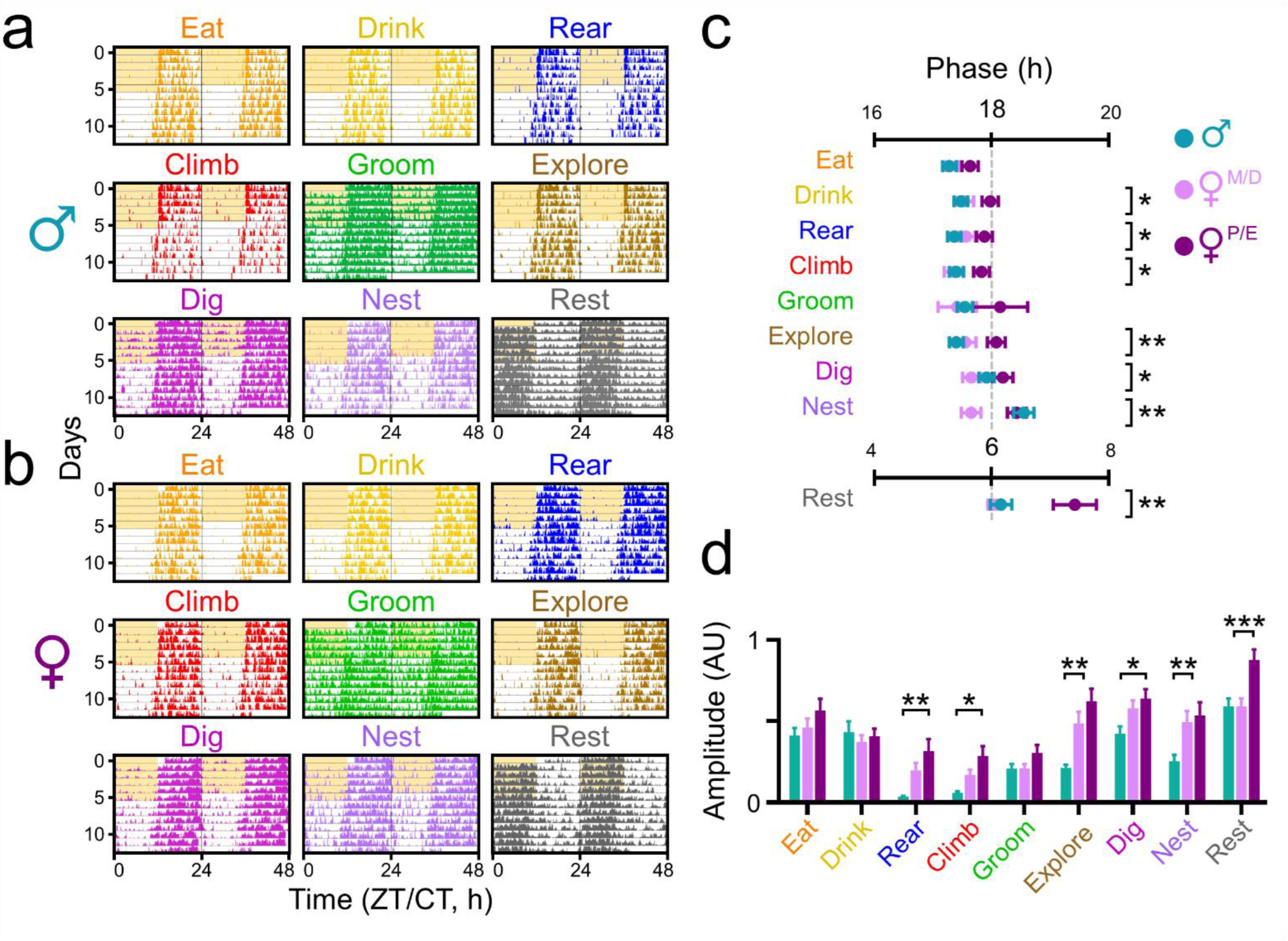
Male and female mice exhibit distinct circadian rhythms in home cage behaviors. **a,b)** Representative double-plotted actograms depicting behaviors (colored lines on each row) averaged across eight male mice or eight female mice that started the experiment in the same estrous state recorded over 5 d in a 12h:12h light:dark (LD) cycle (gray and yellow shading) and 9 d in constant darkness (DD; gray and light gray shading). **c)** Behavior phase comparison plots depicting the acrophases (peak times in circadian time, where CT 18 is subjective midnight and CT 6 is subjective noon) for male (teal, n = 24), metestrus/diestrus (M/D; pink), and proestrus/estrus (P/E; purple) female (n = 27) mice recorded in DD. Lines and error bars depict mean ± SEM. Asterisks indicate behaviors with significant differences in acrophase across groups. One-way ANOVA with post-hoc Tukey’s multiple comparisons test; *, p < 0.05; **, p < 0.01. **d)** Normalized amplitude for each behavior rhythm for male (teal), M/D (pink), and P/E (purple) female mice measured in DD. Two-way ANOVA with post-hoc Tukey’s multiple comparisons test; *, p < 0.05; **, p < 0.01; ***, p < 0.001.

To quantify phase, we measured the acrophase (peak time of activity) for each behavior on each day in LD and in DD (**Fig. 4c; Supplementary Figs. 1a, 2a**). We averaged these acrophases in LD and in DD to more readily compare phase across behaviors and groups. For male mice, we averaged acrophases across each day. We divided female mice into two groups based on their estrous cycle. For “proestrus/estrus” (P/E) female mice, which have relatively high levels of endogenous estrogen, we averaged acrophases over each of the projected days of proestrus based on pre-recording estrous staging. For “metestrus/diestrus” (M/D) female mice, which have relatively low levels of endogenous estrogen, we averaged acrophases over all other days of recording.

Our first goal was to determine if any specific behaviors peaked at distinct times from other behaviors and, if so, whether this pattern was observed in both males and females. To do this, we compared phase markers for all nine behaviors separately within male and female groups (**Supplementary Fig. 2a**). In LD, we found that for all groups of mice, all behaviors except resting and grooming peaked around the same time in the middle of the night (ZT or zeitgeber time 18, where ZT 0 is defined as lights on). As expected for nocturnal animals, resting peaked around the middle of the day, ZT 6. Curiously, grooming behavior in LD in male and P/E (but not M/D) female mice peaked about 30 min and 1.5 h earlier, respectively, than other non-resting behaviors. In DD, for M/D and P/E female mice, all behaviors except resting peaked around the same time, approximately 15 to 30 min earlier than their peak time in LD, as expected for “free-running” nocturnal animals with a shortened period of activity in DD. Surprisingly, for male mice, digging and nesting behaviors were greatly phase delayed in DD. Compared to all other non-resting behaviors, digging and nesting peaked about 30 min and 1 h later, respectively.

Next, to determine if individual behaviors peaked at distinct times in male and female mice, we compared phase markers for each behavior separately across male and female groups. In LD, we found that most behaviors in P/E female mice were phase delayed: eating, drinking, climbing, exploring, and resting behaviors peaked between 30 min to 1 h later compared to these behaviors in M/D females (**Supplementary Fig. 1a**). We also found that in male mice, some behaviors (drinking, grooming, and resting) peaked at similar times to those behaviors in M/D females. However, intriguingly, all other behaviors in male mice (eating, rearing, climbing, exploring, digging, and nesting) peaked at times in between the times those behaviors peaked in M/D and P/E females. In DD, we again found that most behaviors in P/E female mice were phase delayed: drinking, climbing, exploring, digging, nesting, and resting peaked between 30 min to 1 h later compared to these behaviors in M/D females (**Fig. 4c**). Most behaviors in male mice (eating, drinking, rearing, climbing, grooming, exploring, and resting) peaked at similar times to those behaviors in M/D female mice. However, digging and nesting behaviors in male mice instead peaked at similar times to those behaviors in P/E female mice because digging and nesting were phase delayed compared to all other behaviors in male mice in DD. Together, these results demonstrate that behavior rhythms in male and female mice exhibit distinct phase profiles. Specifically, we found that estrous state fundamentally alters behavior phase in female mice and that, in DD, nesting and digging behaviors are significantly delayed in male mice.

We next measured the amplitudes of each behavior rhythm in male, M/D female, and P/E female mice by fitting a cosine wave to their averaged activity profiles in both LD and DD (**Fig. 4d; Supplementary Figs. 1b, 3a**). We found that the amplitudes of all behavior rhythms in male and female mice were dampened in DD compared to LD, consistent with prior reports describing how light cycle influences wheel-running activity amplitude (Li et al., 2006; Pasquali et al., 2010). We also observed that the amplitudes for most behavior rhythms (rearing, climbing, exploring, digging, nesting, and resting) were significantly greater in P/E mice compared to those behaviors in male mice in DD, but not in LD. We found that the amplitudes of some behavior rhythms (climbing, exploring, nesting) were also greater in M/D mice compared to male mice in DD. Finally, we calculated the periods of each behavior rhythm in males and females across all days in both LD and DD (**Supplementary Fig. 4a**). We found that, as expected for nocturnal rodents, the free-running periods in DD for all behavior rhythms in both male and female mice were shorter than the entrained periods in LD (averaged across all behaviors: males LD 24.02 ± 0.03 h; males DD 23.73 ± 0.04 h; females LD 24.04 ± 0.03 h; females DD 23.82 ± 0.03 h). Surprisingly, sex had little effect on period. Digging in females exhibited a slightly lengthened period in DD, but no other behaviors showed significant period differences. Together, these results demonstrate that biological sex has a profound effect on the amplitude of most behavioral rhythms but has little to no effect on periodicity.

### Estrogen replacement phenocopies multiple behavior rhythm changes seen in proestrus female mice

To determine whether these observed sex differences in circadian behavior could be explained by differences in endogenous estrogen levels, we again continuously recorded videos, inferred behaviors, and generated actograms from ovariectomized (OVX; n = 24) and ovariectomized, estradiol-supplemented (OVXE; n = 22) female mice over 5 d in LD and over 5 d in DD (**Figs. 5a, b**) (Ström et al., 2012). OVX mice have chronically low levels of estrogen similar to the levels found in male mice or during metestrus/diestrus in intact females, and OVXE mice have chronically elevated levels of estrogen similar to the levels found during proestrus in intact females. We used these actograms to again determine key circadian properties of each rhythm including phase, amplitude, and period.

**Figure 5.**
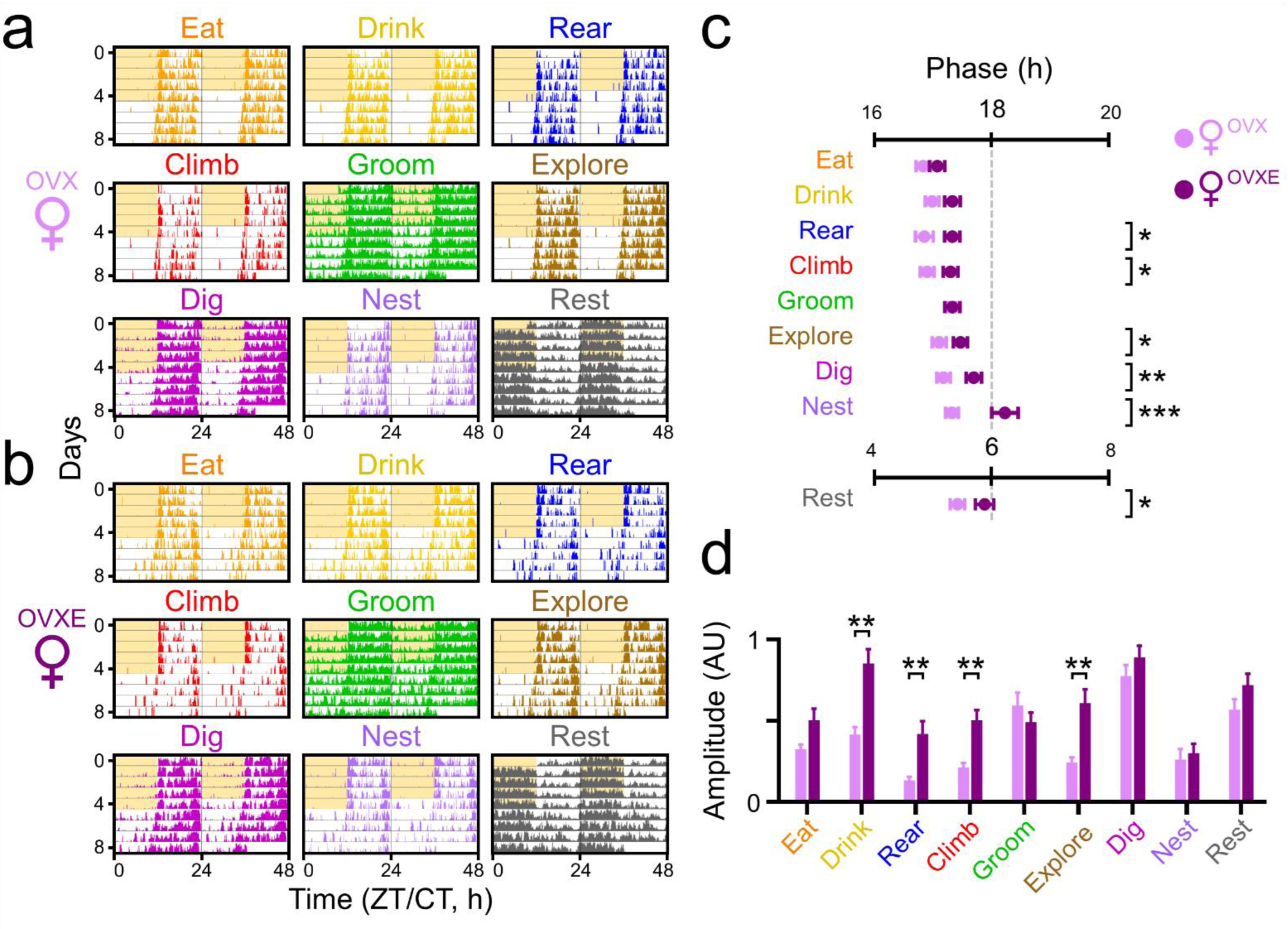
Ovariectomized and ovariectomized, estradiol-supplemented female mice exhibit distinct circadian rhythms in home cage behaviors. **a,b)** Representative double-plotted actograms depicting behaviors (colored lines on each row) averaged across eight ovariectomized (OVX) female mice or eight ovariectomized, estradiol-supplemented (OVXE) female mice recorded over 5 d in a 12h:12h light:dark (LD) cycle (gray and yellow shading) and 5 d in constant darkness (DD; gray and light gray shading). **c)** Behavior phase comparison plots depicting the acrophases (peak times in circadian time, where CT 18 is subjective midnight and CT 6 is subjective noon) for OVX (pink, n = 24) and OVXE (purple; n = 22) female mice recorded in DD. Lines and error bars depict mean ± SEM. Asterisks indicate behaviors with significant differences in acrophase across groups. One-way ANOVA with post-hoc Tukey’s multiple comparisons test; *, p < 0.05; **, p < 0.01. **d)** Normalized amplitude for each behavior rhythm for OVX (pink) and OVXE (purple) female mice measured in DD. Two-way ANOVA with post-hoc Tukey’s multiple comparisons test; **, p < 0.01.

To quantify phase, we again measured the acrophase (peak time of activity) for each behavior on each day in OVX and OVXE mice in LD and in DD (**Fig. 5c; Supplementary Figs. 1c, 2b**). We averaged these acrophases in LD and in DD to more readily compare phase across behaviors and groups. First, to determine if any specific behaviors peaked at distinct times from other behaviors in OVX and OVXE mice, we compared phase markers for all nine behaviors separately within estrogen replacement groups (**Supplementary Fig. 2b**). In LD, we found that for both OVX and OVXE mice, all behaviors except resting peaked around the same time in the middle of the night (ZT 18); resting peaked around the middle of the day, ZT 6. In DD, for both OVX and OVXE mice, most non-resting behaviors peaked around the same time, about 30 min earlier than their peak time in LD as expected for nocturnal rodents. However, grooming in OVX mice peaked about 30 min later, and nesting and resting in OVXE mice peaked about 30 min and 1 h later, respectively, compared to all other non-resting behaviors.

Next, to determine if individual behaviors peaked at distinct times in OVX and OVXE mice, we compared phase markers for each behavior separately across estrogen replacement groups. In LD, we found no difference between OVX and OVXE mice in the peak times of any behavior (**Supplementary Fig. 1c**). In DD, similar to what we observed with intact P/E female mice, most behaviors (rearing, climbing, exploring, digging, nesting, and resting) in OVXE mice were phase delayed, peaking between 30 min and 1 h later compared to these behaviors in OVX mice (**Fig. 5c**). However, surprisingly, eating, drinking, and grooming behaviors in OVXE mice peaked at the same time as those behaviors in OVX mice. Together, these results demonstrate that behavior rhythms in OVX and OVXE mice exhibit distinct phase profiles. Specifically, we found that estrogen replacement significantly phase delays most, but, importantly, not all behaviors in DD, but not in LD.

We next measured the amplitudes of each behavior rhythm in OVX and OVXE mice by fitting a cosine wave to their averaged activity profiles in both LD and DD (**Fig. 5d; Supplementary Figs. 1d, 3b**). We found that the amplitudes of all behavior rhythms except eating and nesting in OVX and OVXE mice were dampened in DD compared to LD. We also observed that the amplitudes for some, but not all, behavior rhythms (drinking, rearing, climbing, and exploring) were significantly greater in OVXE mice compared to those behaviors in OVX mice in DD, but not in LD. Finally, we calculated the periods of each behavior rhythm in OVX and OVXE mice across all days in both LD and DD (**Supplementary Fig. 4b**). We found that, as expected, the free-running periods in DD for all behavior rhythms but nesting in both OVX and OVXE mice were shorter than the entrained periods in LD (averaged across all behaviors: OVX LD 24.07 ± 0.02 h; OVX DD 23.70 ± 0.06 h; OVXE LD 24.04 ± 0.04 h; OVXE DD 23.66 ± 0.06 h). Estrogen replacement had little effect on period: only drinking and grooming behaviors in OVXE mice had slightly different periods (longer and shorter, respectively) than in OVX mice. Together, these results demonstrate that estrogen replacement greatly influences both the phase and amplitude of multiple behavior rhythms. Specifically, estrogen replacement increases behavior rhythm amplitudes and mimics the phase delays we observed in intact female mice during proestrus.

### A generalizable circadian behavioral analysis suite

Our DINOv2+ model and automatic inference system allowed us to thoroughly investigate how circadian behaviors are influenced by sex and estrogen levels at an unprecedented throughput and acquisition rate. We realized that introducing our software infrastructure to the broader scientific community could be revolutionary to fields seeking to understand the temporal characteristics of animal behavior, including, particularly, circadian biology. We therefore developed a “circadian behavioral analysis suite” (CBAS), a Python package aimed at generalizing the software and DINOv2+ model used in our experiments (**Fig. 6a**). CBAS is equipped to handle automated, continuous video acquisition, automated inference using the DINOv2 feature extractor and joint long short-term memory (LSTM) and linear layer models, and visualization of behavior actograms in real time (**Fig. 6b**). Briefly, CBAS is divided into three modules: an acquisition module, a classification and visualization module, and an optional training module. The acquisition module is capable of batch processing streaming video data from any number of network-configured real-time streaming protocol (RTSP) IP cameras. The classification and visualization module enables real-time inference on streaming video and displays acquired behavior time series data in real time as actograms that can be readily exported for offline analysis in a file format compatible with ClockLab Analysis, a widely-used circadian analysis software. Users wanting to fully replicate our recording setup (see the Jones Lab Github for a full parts list and assembly instructions) and nine behaviors of interest can immediately begin classification using our DINOv2+ joint LSTM and linear layer model that is included in the Python package. Importantly, because the DINOv2 visual backbone is kept static in our training model, users can also quickly and easily adapt CBAS to accommodate a diverse array of classification tasks, animal species, and video environments. The training module allows the user to create balanced training sets of behaviors of interest, train joint LSTM and linear layer model heads, and validate model performance on a naive test set of behavior instances. Importantly, CBAS’s acquisition module is essentially machine learning model agnostic, allowing for future models to be easily incorporated into the CBAS pipeline. Together, these modules present an intuitive, accessible software interface that will allow for the rapid adoption of CBAS by end users with any level of programming ability.

**Figure 6.**
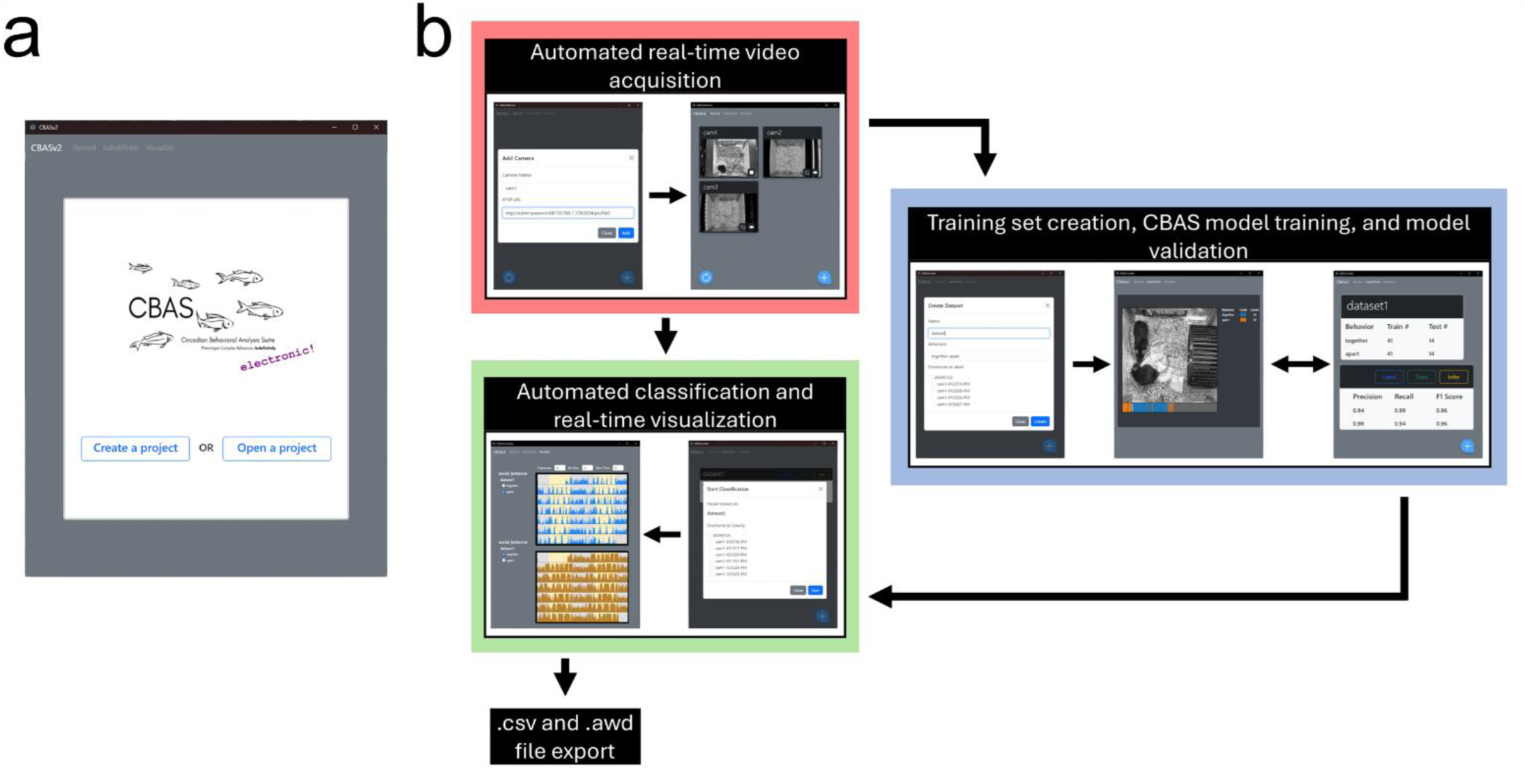
CBAS: a circadian behavioral analysis suite. **a)** CBAS is a user-friendly GUI-enabled Python package that allows for the automated acquisition, classification, and visualization of behaviors over time. **b)** Schematic of the CBAS pipeline. Red; acquisition module; blue, training module; green, classification and visualization classification module.

## Discussion

Here, we developed and validated a novel system for transforming existing machine learning classifiers into real-time sensors capable of phenotyping circadian rhythms in complex behaviors for an indefinite length of time. We used this pipeline to thoroughly characterize the effects of biological sex and estrogen levels on circadian behavior across 97 individual mouse recordings with a minimum duration of 10 d per recording at 10 frames per second. We then developed this toolkit into an open-source, user-friendly Python package – CBAS – for use by the broader circadian biology community and beyond. CBAS has the potential to reveal temporal variations in behavior that have previously gone undetected in a diverse range of animal models. In addition, CBAS provides scientists with the tools needed to build, adequately validate, and automate highly reliable machine learning classifiers for any complex behavior(s) of interest.

CBAS’s extensive model validation, classification power, and customizable, open-source nature set it apart from previous commercial (e.g., HomeCageScan, (Adamah-Biassi et al., 2013, 2014)) and non-commercial (e.g., (Steele et al., 2007; Goulding et al., 2008; Jhuang et al., 2010; Salem et al., 2015)) home cage behavior acquisition tools. For instance, (Jhuang et al., 2010) uses background masking and motion features to classify behaviors with a Hidden Markov Model Support Vector Machine, an outdated, but theoretically capable, architecture. While the authors point to a high classification accuracy, there is little to no information provided regarding more standard machine learning classification metrics such as precision, recall, F1 score, etc. With some assumptions, some of these values can be calculated from the provided confusion matrices, but for most behaviors they fail to meet our strict model performance threshold. Critically, the class balance of the training and test sets used for their model verification is unreported, and neither set remains available to the public. Similarly, (Goulding et al., 2008) fails to perform model performance validation for their supervised learning technique, although they do show that their system is capable of recording behaviors on a circadian timescale. However, because their model only identifies active and inactive behavior states, it is incapable of automatically identifying complex behaviors or animal-environment interactions. Adequate performance metrics, training sets, and testing sets of the HomeCageScan system are likewise scarce, a problem that is exacerbated by its closed-source nature. Importantly, none of these systems has user-friendly customizability to extend classification to other animal models, environments, and novel behaviors. Our goal with CBAS is to allow users with any level of programming ability to integrate completely customizable machine learning models, extensively validate model performance, and record behaviors in real time, indefinitely.

Although there have been several recent advances in supervised and even unsupervised pose-based behavior classification (e.g., SimBA, B-SOiD, A-SOiD, (Nilsson et al., 2020; Hsu and Yttri, 2021; Tillmann et al., 2024)), pose-based classifiers sacrifice critical learnable information with their sweeping dimensional reduction to pose dynamics. For example, the subtle home cage environment differences that characterize digging versus nesting behavior in our recording setup would be completely lost in a reduction to pose time series. Furthermore, labeling pose data is significantly more difficult than labeling classification data, especially in dynamic video environments or with moving subjects. In contrast, DeepEthogram (DEG) models and our proposed DINOv2+ model have the capacity to learn specific high-dimensional features from spatial and temporal dynamics derived from raw pixel values (Bohnslav et al., 2021). Importantly, DEG models and DINOv2 are fundamentally different in how they are trained. DEG, which we previously used to analyze circadian behavior in a proof-of-concept study (Wahba et al., 2022), uses a feature extractor that is trained in a supervised manner where the model receives direct classification feedback from labeled data throughout training. In contrast, the DINOv2 feature extractor is pretrained in a self-supervised manner where the model is encouraged to produce a rich, often clustered, visual feature space that can then be used as a frozen basis for subsequent small supervised classification models. The DINOv2 backbone model has several benefits because of its self-supervised learning strategy. Most notably, this strategy generalizes the model’s feature space to data and tasks which it has never been trained to recognize or accomplish. For example, DINOv2 is broadly capable of semantic segmentation, depth estimation, image/video classification, or object tracking/recognition with the minor addition of a trainable linear network layer (Oquab et al., 2023). Reducing bias is closely linked to enhancing feature generalizability. While supervised learning often optimizes for unstable visual heuristics, self-supervised training disregards these heuristics in favor of robust visual characteristics (Caron et al., 2021; Shwartz-Ziv and LeCun, 2023). Finally, self-supervised models help bridge the growing gap in access to computing resources and data science expertise needed to fully train and optimize high-performing vision models.

In addition to using our DINOv2+ model for inference, CBAS also leverages our standardized video recording pipeline to allow for real-time behavior classification that can rival the throughput of sensor-based recording systems (Siepka and Takahashi, 2005; Verwey et al., 2013). CBAS is also incredibly cost effective. The total cost of the hardware we used in this study, including our custom-built mouse cages and circadian behavior cabinets, is only two-thirds the estimated cost of standard commercial systems. Moreover, our setup can be used to record and infer any number of behaviors in parallel, whereas standard circadian acquisition hardware is only capable of recording locomotor behavior by measuring wheel-running activity or infrared beam breaks. Another major advantage of CBAS is its accessibility, presenting a low barrier to entry for the broader circadian research community, including those with limited programming skills. Additionally, CBAS mirrors the functionality of field-standard circadian analysis systems by plotting behavior data as actograms in real time, providing immediate feedback about the state of ongoing experiments. CBAS also outputs these behavior actograms in formats compatible with Actimetrics’ ClockLab Analysis software, which ensures that researchers can adapt familiar analyses to CBAS-generated behavior data. The open-source nature of CBAS essentially democratizes the circadian analysis of complex behaviors, allowing a greater number of researchers to investigate long-term behavior dynamics.

Previous studies have identified numerous sex differences in the temporal patterning of physiology. For example, male and female mice exhibit distinct circadian rhythms in glucocorticoid production, cardiovascular function, body temperature, and immune function (Griffin and Whitacre, 1991; Atkinson and Waddell, 1997; Sanchez-Alavez et al., 2011; Barsha et al., 2016; Walton et al., 2022). However, the question of whether there are pronounced sex differences in the temporal patterning of behavior has, to date, been mostly unanswered. Subtle sex differences have been observed in wheel-running activity rhythms (Lee et al., 2004; Kuljis et al., 2016; Anderson et al., 2023). For instance, male mice show a greater precision of wheel-running activity onsets in LD and female mice show a longer wheel-running activity duration on the day of proestrus in DD (Albers et al., 1981; Kuljis et al., 2013). However, given the presumed global regulation by the circadian system of multiple brain circuits that control distinct behaviors, more work is needed to reveal and understand sex differences in other behavioral rhythms (Starnes and Jones, 2023). We used CBAS to address this by testing the hypothesis that circadian rhythms in behavior differ between males and females. We identified differences in behavioral rhythms including, notably, that nesting and digging rhythms exhibit a distinct phase delay in male, but not female, mice during constant darkness that had not been previously reported. We also observed that most behavioral rhythms in female mice had higher peak-to-trough amplitudes, which is suggestive of more robust circadian organization. This is consistent with previous work showing that the amplitude of wheel-running activity rhythms is greater in female mice compared to male mice (Anderson et al., 2023). Previous studies have also identified that the duration of wheel-running activity is extended (that is, it ends later) on the day of proestrus (Albers et al., 1981). We confirm this finding and extend it to demonstrate that the temporal organization of nearly all behaviors changes across the estrous cycle. Our findings reveal critical unseen sex differences in many, but, importantly, not all behavioral rhythms, which emphasizes the importance of measuring circadian rhythms in behaviors other than locomotor activity.

The limited number of previously identified sex differences in circadian behavior have been speculated to be due to differences in levels of circulating sex hormones and/or sex hormone receptor expression (Walton et al., 2022). Our experiments were designed to allow us to distinguish between differences in behavioral rhythms that are due to biological sex and those that are due to the presence or absence of estrogen. We found that estrogen replacement recapitulates most, but not all, of our observed sex differences in behavioral rhythms. For instance, we found that P/E females exhibit higher amplitude circadian rhythms in most behaviors compared to males and M/D females. Similarly, many behavioral rhythms in OVXE females are more robust than in OVX females. We also found that the peak time of most behaviors in “high estrogen” P/E and OVXE females was delayed compared to “low estrogen” M/D and OVX females. One possible explanation for this is the relative distribution of estrogen receptors in brain circuits that regulate different behaviors. Exogenous estrogen has been shown to increase the amplitude of wheel-running activity rhythms through the activation of estrogen receptor (ER)α but to delay the phase of wheel-running activity rhythms through the activation of ERβ (Royston et al., 2014). Indeed, a recent study determined that the lateral hypothalamus (LH), which has subpopulations of both ERα-positive and ERβ-positive neurons, regulates nest-building behavior (Merchenthaler et al., 2004; Sotelo et al., 2022). If these ERβ-expressing LH neurons are preferentially activated during nesting behavior, this could explain why estradiol delays nesting rhythms in OVXE and P/E mice. However, this does not explain why male mice, which have low levels of endogenous estrogen, exhibit delayed digging and nesting rhythms that peak at similar times to those rhythms in OVXE and P/E mice, which have high levels of endogenous estrogen. Further studies will need to determine whether this finding is due to sex differences in developmental circuit wiring, differences in estrogen receptor distribution and/or expression levels, or other factors.

In this study, we used our circadian behavioral analysis suite (CBAS) to automatically quantify differences in the circadian regulation of behavior between male and female, and OVX and OVXE female, mice. This approach can be readily expanded to address other critical questions in circadian biology, neuroscience, and ecology, including the ethological investigation of other behavioral rhythms in videos of mice recorded in the laboratory and, potentially, in the wild. Notably, CBAS can also be used for the rapid circadian phenotyping of mice with different genotypes or disorders (Richardson, 2015). Current approaches almost universally measure changes to wheel-running activity rhythms as evidence that a mutation, gene, or drug influences circadian behavior. Here, we found that some, but, critically, not all, behavioral rhythms differ by biological sex and by estrogen levels. It is therefore highly likely that any given experimental treatment could cause circadian alterations in behaviors other than, or in addition to, wheel-running activity. CBAS aims to extend the modern toolkit of machine learning classification into any and all long-term behavior assays, greatly expanding the scope of potential hypotheses and impact of future studies.

## Materials and methods

### Animals

Prior to recording, we group-housed male (n = 24) and female (n = 51; 24 of which were subsequently ovariectomized, see next section) wild-type mice in their home cages in a 12h:12h light:dark cycle (LD, where lights on is defined as zeitgeber time (ZT) 0; light intensity ∼2 x 10^14^ photons/cm^2^/s) at constant temperature (∼23°C) and humidity (∼40%) with food and water provided ad libitum. All mice were between 6 and 12 weeks old at the time of the recording. To determine the estrous stage of female mice, we performed vaginal lavage for four consecutive days prior to beginning long-term recording (Byers et al., 2012). All experiments were approved by and performed in accordance with the guidelines of Texas A&M University’s Institutional Animal Care and Use Committee.

### Ovariectomy and estradiol capsule implantation

We ovariectomized a cohort of female mice (OVX, n = 24) using standard methods (Ström et al., 2012). Briefly, we made a sterile ∼2 cm bilateral incision through the skin and peritoneum immediately dorsal to the ovaries. After ligating and removing each ovary, we sutured the peritoneum and skin incisions. We provided the mice with buprenorphine-SR (1 mg/kg; subcutaneous) and enrofloxacin (0.25 mg/ml; ad libitum in their drinking water) and allowed them to recover in their home cages for at least 1 week prior to beginning long-term recording. After recording OVX mice, we implanted them with a sterile 2 cm silastic capsule containing 17-β-estradiol (36 μg/ml in sesame oil) subcutaneously between the shoulder blades (OVXE; n = 22) (Ström et al., 2012). We excluded two OVX mice from capsule implantation because they had excessive barbering around their ovariectomy incision site. We allowed OVXE mice to recover in their recording cages for 1 d prior to beginning long-term recording.

### Experimental housing

We transferred individual mice from their home cages to custom-built recording cages inside custom-built light-tight, temperature-and humidity controlled circadian cabinets for the duration of our experiments. We built the cages (external dimensions, length x width x height: 22.9 cm x 20.3 cm x 21.6 cm; internal dimensions: 20.3 cm x 17.8 cm x 20.3 cm) out of transparent and opaque acrylic panels (thickness, walls and floor: 6.4 mm; lid, 3.2 mm) and T-slot aluminum extrusions (25.4 mm^2^) (**Fig. 1a**). We 3D printed custom water bottle holders and food hoppers out of PLA filament, coated them in food safe clear-cast epoxy resin (Alumilite), and affixed them to the acrylic walls. To continue our recordings throughout the dark phase, and to prevent potential glare and shadows from ceiling-mounted lights, we affixed dim infrared (850 nm) light strips to the cage lid using a custom 3D printed cage topper. Prior to recording, we added ∼7 mm wood chip bedding and a 25 mm by 50 mm square of cotton nestlet to the cage bottom, added food pellets to the food hopper, and attached a standard water bottle filled with water to the water bottle holder such that its metal spout protruded about 1.5 cm into the cage. Our circadian cabinets were built to hold twelve of our custom-built mouse cages across three vertical shelves, with four cages per shelf. We controlled the ceiling-mounted lights in the cabinets (broad-spectrum white light, ∼6 x 10^13^ photons/cm^2^/s measured at the cage floor) using ClockLab Data Collection hardware and software (Actimetrics) that communicated via a 5V transistor-transistor logic signal with a high-power power relay (Digital Loggers). We performed daily animal welfare checks using dim red light (650 nm).

### Automated video recording

We positioned power-over-internet (PoE) IP cameras without infrared filters (I706-POE, Revotech) equipped with 6 mm lenses (Xenocam) 47.5 cm above the recording cages such that all four corners of the cage, the food hopper, and the water spout were each visible in the recorded video and the mouse and nesting material were in focus. We recorded all videos at 10 frames per second (fps) with in-camera image settings set to a contrast of 130/255, brightness of 140/255, saturation of 0/255, and sharpness of 128/255. We streamed videos at a main stream bitrate of 2048 kilobits per second (kb/s) and a secondary stream bitrate of 256 kb/s. We disabled audio streams to reduce bandwidth. We recorded our mice in cohorts of 8 to 12 mice split between two circadian cabinets. We paused our recordings briefly between light settings (LD and DD) to allow for cage changes, if necessary.

We used FFmpeg, a standard open-source video processing tool, to develop a custom video acquisition system capable of streaming live video and storing successively binned segments of video for each network camera simultaneously. During a recording, FFmpeg automatically handles cropping the video to a region of interest and scaling the video to a desired size (here to a scale of 256×256 pixels). To do this, we connected our cameras in parallel via 10 gigabits per second (Gbps) Cat6 ethernet cables to a Gigabit PoE switch (Aruba JL684A#ABB). We then connected our switches via 40 Gbps Cat8 ethernet cables to our custom machine-learning computers (12-core AMD Ryzen 9 5900X CPU, 32 GB RAM, NVIDIA GeForce RTX 3090 with 24 GB VRAM) (**Fig. 3a**). During a recording, our software creates two threads to monitor the creation of streamed video segments and orchestrate the inference of incoming data (the “storage” and “inference” threads). The storage thread records information about each video segment (creation time, segment length, camera-specific settings), moves the video segments to the corresponding camera directories on the computers, and notifies the inference thread that new videos are available for inference (see below).

### Behavior definitions

We defined a list of nine home cage behaviors (eating, drinking, rearing, climbing, grooming, exploring, digging, nesting, and resting, (Garner, 2017)) with the goal of identifying the visual and motion characteristics of each behavior that our DINOv2+ model would be capable of learning (**Supplementary Table 1**). As such, our definitions do not aim to ascribe intent to a behavior (as humans are often inclined to do), but rather contain references to particular features that strictly define behavioral classes. These include spatial features that are necessary constraints on a behavior and temporal features that are split into two groups indicative of the start and stop of a behavior sequence. To further enforce these rigorous criteria defining behaviors, our entire set of training instances were generated by a single labeler.

### Model training and inference

To train our baseline DeepEthogram (DEG) classifier, we needed to individually train three components, a “flow generator” that estimates optic flow across video frames, a “feature extractor” that determines the probability of a behavior being present on a given frame based on a low-dimensional set of temporal and spatial features, and a “sequence model” that further refines model predictions using a temporal gaussian mixture (TGM) model with a larger temporal receptive field. We trained our flow generator on a set of videos consisting of approximately 500,000 frames of videos from 8 mice recorded at 10 fps. We then trained our feature extractor using the medium model size preset (deg_m, (Bohnslav et al., 2021)) and our TGM sequence model using a temporal window of 15 frames. We trained both the feature extractor and TGM sequence models on an identical balanced training set used for subsequent training of our DINOv2+ model (see below). Importantly, we include our model configuration files for all DEG models, our DINOv2+ model, and all trained model weights at the Jones lab Google Drive repository (see Data Availability section below).

Our DINOv2+ model architecture was designed to take as input sequenced outputs from the DINOv2 feature extractor and produce a robust, frame-to-frame stable, and accurate classification time series (**Fig. 2a**). The DINOv2 feature extractor model outputs one 768 length vector for each given video frame encoding the relevant visual information about the image scene. Our joint long short-term memory (LSTM) and linear layer classification head integrates visual and motion information from a sequence of vectors (here 31 frames) centered at the frame of interest into a behavioral classification. During a forward pass of our classification head, noise is randomly injected into a normalized version of the sequence of DINOv2 outputs, transformed through a single linear layer into the output size, and then averaged over an 11 frame, centered sub-window. Simultaneously, the mean of the original input sequence is subtracted from the input sequence, compressed by a linear layer to a latent dimension, and then passed through a single layer bidirectional LSTM network. The logits of the LSTM layer are condensed to the output size and added to the outputs of the linear layer. A softmax of the summed output results in the model’s behavior classification confidence for each frame. The softmax function is defined as

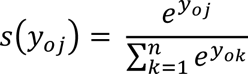

where *n* is vector length (here, 9), *y*_*oj*_is the output vector at position *j*, and *y*_*ij*_ is the input vector at position *j*.

We next trained DINOv2+ classification head on a balanced set of behavior instances sampled across the light and dark phases from 30 unique mice and cages. For this task, we trained our model using a cross entropy loss function, defined as:

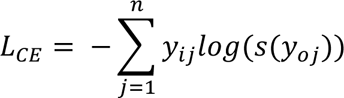

where, again, *n* is vector length (here, 9), *y*_*oj*_is the output vector at position *j*, and *y*_*ij*_ is the input vector at position *j* (Ciampiconi et al., 2023). Additionally, we added a covariance loss to discourage covariance of our LSTM output features. Our covariance loss was defined as the off-diagonal sum of the absolute covariance matrix constructed using the raw latent dimensional outputs of the LSTM layer divided by our latent dimension size. This approach was inspired by the elegant loss function employed in the VICReg learning scheme, and it consistently improved our classification performance <u>(Bardes et al. 2021)</u>. We identified optimal hyperparameters that minimized the total loss to be a latent dimension of 256, an LSTM latent dimension of 64, and a linearly decreased learning rate of 5e-4 to 1e-5 over 10 epochs of training. During classification training, model states are selectively saved by maximizing for the weighted average F1 score of model performance on a test set.

### Model validation

To validate the performance of our behavior classifier, we used a naive, balanced test set of behavior sequences. Prior to model training, we randomly selected each unique behavior sequence (or “instance”) from our annotated dataset while preserving class balance. Importantly, to prevent misleading or skewed model performance results, we did not use the instances in this test set during any form of model training or adjustment.

From this balanced test set, we randomly sampled 1,000 sequences with a maximum length of 31 frames. After we used our model to infer all sampled sequences, we calculated precision, recall, F1 score, specificity, and balanced accuracy using the *sklearn.metrics* library in Python. We also calculated the normalized Matthews correlation coefficient (nMCC) using a custom Python implementation (Chicco and Jurman, 2020). We repeated this random sampling for a total of ten iterations before calculating the mean and standard deviations of each metric.

To cross validate our DINOv2+ model with the DEG TGM sequence model and human annotators, we first trained a TGM sequence model with a temporal window of 15 frames on the equivalent training set to that of our DINOv2+ model. We then repeated the sampling and metric calculation detailed above to determine means and standard deviations for the TGM model metrics. Using both models’ output prediction probabilities, we calculated the precision-recall curves for each classifier. To determine if the change between the area under the precision-recall curves (AUPRC) was significant, we used a Python version of a bootstrapping method originally implemented in R to create a normal distribution of area differences between random subsamples of the two curves with 10,000 sampling iterations (Zobolas, n.d.). We then compared the area difference of the two total precision-recall curves to the mean and standard deviation of this distribution to determine significance.

Finally, we designed a custom GUI in Python that allows human annotators to classify 10,000 randomly sampled 15 frame sequences of video frames from our test set by replaying the sequence until it is classified as a behavior. Importantly, the GUI does not give the human annotator performance feedback over the course of the annotation so (much like our machine learning models) they are unable to learn as they annotate. Using these annotations, we repeated the sampling and metric calculations to determine means and standard deviations for human labeler metrics.

### Automated inference

At the beginning of a recording, our software automatically bins the video stream into segments of time. In these experiments, we chose to record in thirty minute intervals. For each new video bin, a subprocess infers the video using the frozen DINOv2 feature extractor model. In this manner, CBAS continuously automates model inference until the recording is terminated by the user. Users can also add pre-recorded videos to the project directory to begin the DINOv2 inference of these videos. If a joint LSTM and linear layer model is trained and ready for use in inference (as in our experiments), CBAS also coordinates the automated inference of the DINOv2 features into sequenced behavior classes.

### Analysis

We produced behavior actograms by binning the number of frames predicted as a given behavior over a 30 min period. To account for differing estrous states in female mice, we shifted their behavior actograms such that their projected day of proestrus (as determined by estrous scoring) was aligned for each mouse that we recorded. In LD, adjustments and group ns were: 0 d (n = 8 mice), -1 d (n = 5 mice), -2 d (n = 7 mice), and -3 d (n = 7 mice). In DD, adjustments and group ns were: 0 d (n = 5 mice), -1 d (n = 4 mice), -2 d (n = 6 mice), -3 d (n = 13 mice).

To calculate circadian parameters (phase, period, and amplitude), we used CBAS to export each actogram as an .awd file, a file format compatible with ClockLab Analysis (Actimetrics), a widely-used circadian analysis software. To calculate phase, we identified acrophases by calculating the midpoint between onset and offset times determined by a standard template matching algorithm that searches for a 12 h period of inactivity (or activity) followed by a 12 h period of activity (or inactivity). For analysis, acrophases for male, OVX, and OVXE mice were averaged across each day in LD and DD. Acrophases for proestrus/estrus (P/E) female mice were averaged on the projected days of proestrus: days 1 and 5 in LD and days 1, 5, and 9 in DD. Acrophases for metestrus/diestrus (M/D) female mice were averaged on all other days (days 2-4 in LD and days 2-4 and 6-8 in DD). To calculate period, we used a Lomb-Scargle periodogram with a range of 20 to 28 hours and a significance level of 0.001. To calculate amplitude, we measured the peak-to-peak amplitude of a sine wave fitted to the average activity profile calculated across all days in LD or all days in DD.

We performed the following statistical tests in Prism 10.0 (Graphpad): one-way ANOVA, unpaired t-test, two-way ANOVA, Tukey’s multiple comparisons test, Dunnett’s multiple comparisons test. We performed a bootstrapping test in Python (Zobolas, n.d.). Because no phase markers occurred at or near the 24 h modulus, we performed statistical comparisons without using circular statistics. We used Shapiro-Wilk and Brown-Forsythe tests to test for normality and equal variance, defined α as 0.05, and presented all data as mean ± SEM.

## Data availability

All data generated in this study that support our findings are presented within this paper or its Supplementary Materials or at the Jones lab Google Drive repository at http://tinyurl.com/jones-lab-tamu. CBAS is also available to the public at the Jones lab Github page at https://github.com/jones-lab-tamu.

## Acknowledgments

We thank the members of the Jones lab for discussion and comments on the manuscript and V. Fisher for assistance with ovariectomies. This work was supported by National Institutes of Health Grant R35GM151020 (J.R.J.) and a Research Grant from the Whitehall Foundation (J.R.J.).

