## Supplementary Figures for "A circadian behavioral analysis suite for real-time classification of daily rhythms in complex behaviors"

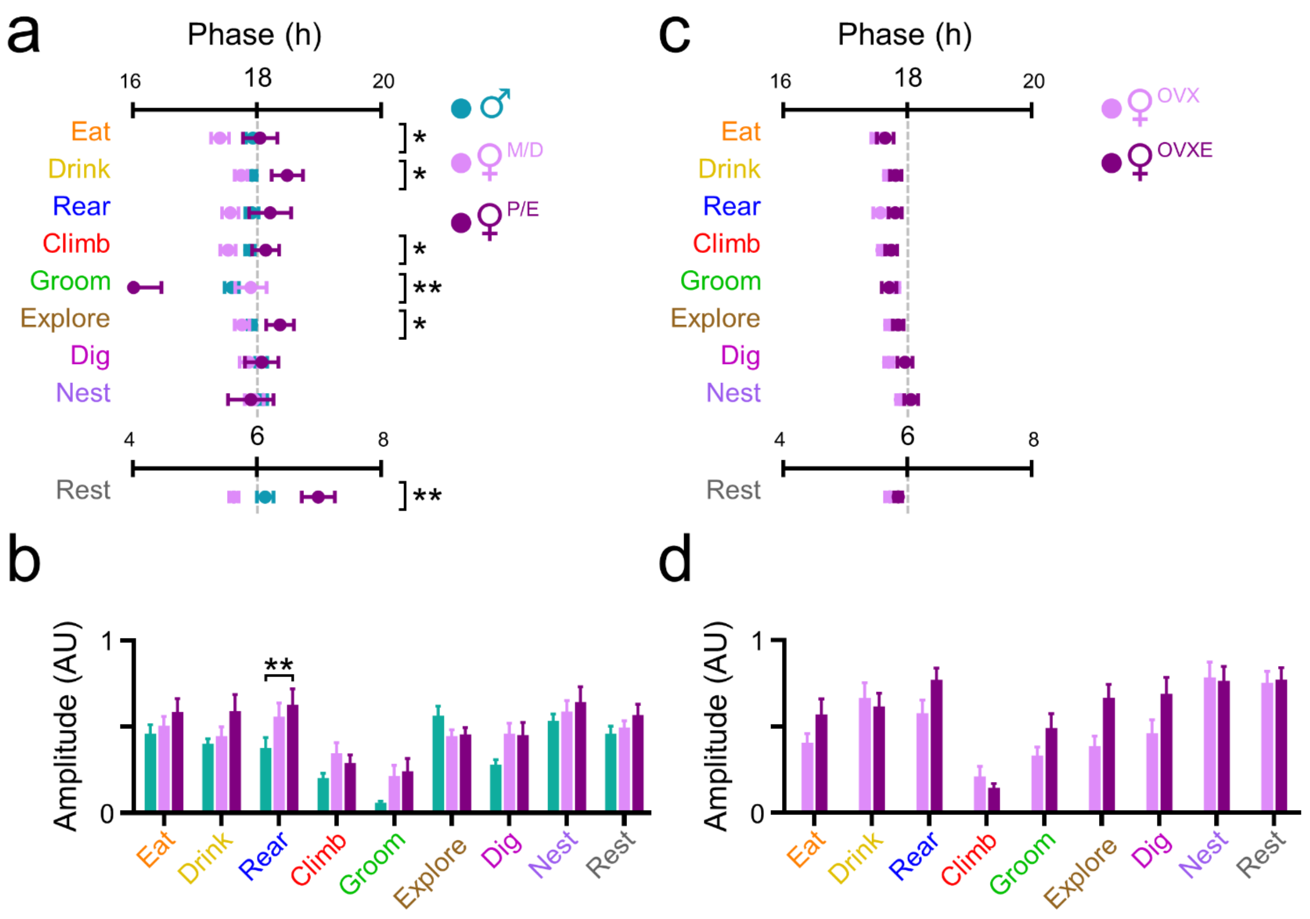


**Supplementary Figure 1. Sex and estrogen effects on behavior phase and amplitude in LD. a,c)** Behavior phase comparison plots depicting the acrophases (peak times in zeitgeber time, where ZT 18 is midnight and ZT 6 is noon) for **(a)** male (teal, n = 24), metestrus/diestrus (M/D; pink), and proestrus/estrus (P/E; purple) female (n = 27) mice, or **(c)** ovariectomized (OVX; pink; n = 24) and ovariectomized, estradiol-supplemented (OVXE; purple, n = 22) female mice, recorded in a 12h:12h light:dark (LD) cycle. Lines and error bars depict mean ± SEM. Asterisks indicate behaviors with significant differences in acrophase across groups. One-way ANOVA with post-hoc Tukey's multiple comparisons test; *, $p < 0.05$; **, $p < 0.01$. **b,d)** Normalized amplitude for each behavior rhythm for **(a)** male (teal), M/D (pink), and P/E (purple) female mice or **(d)** OVX (pink) and OVXE (purple) female mice measured in LD. Two-way ANOVA with post-hoc Tukey's multiple comparison's test; **, $p < 0.01$.

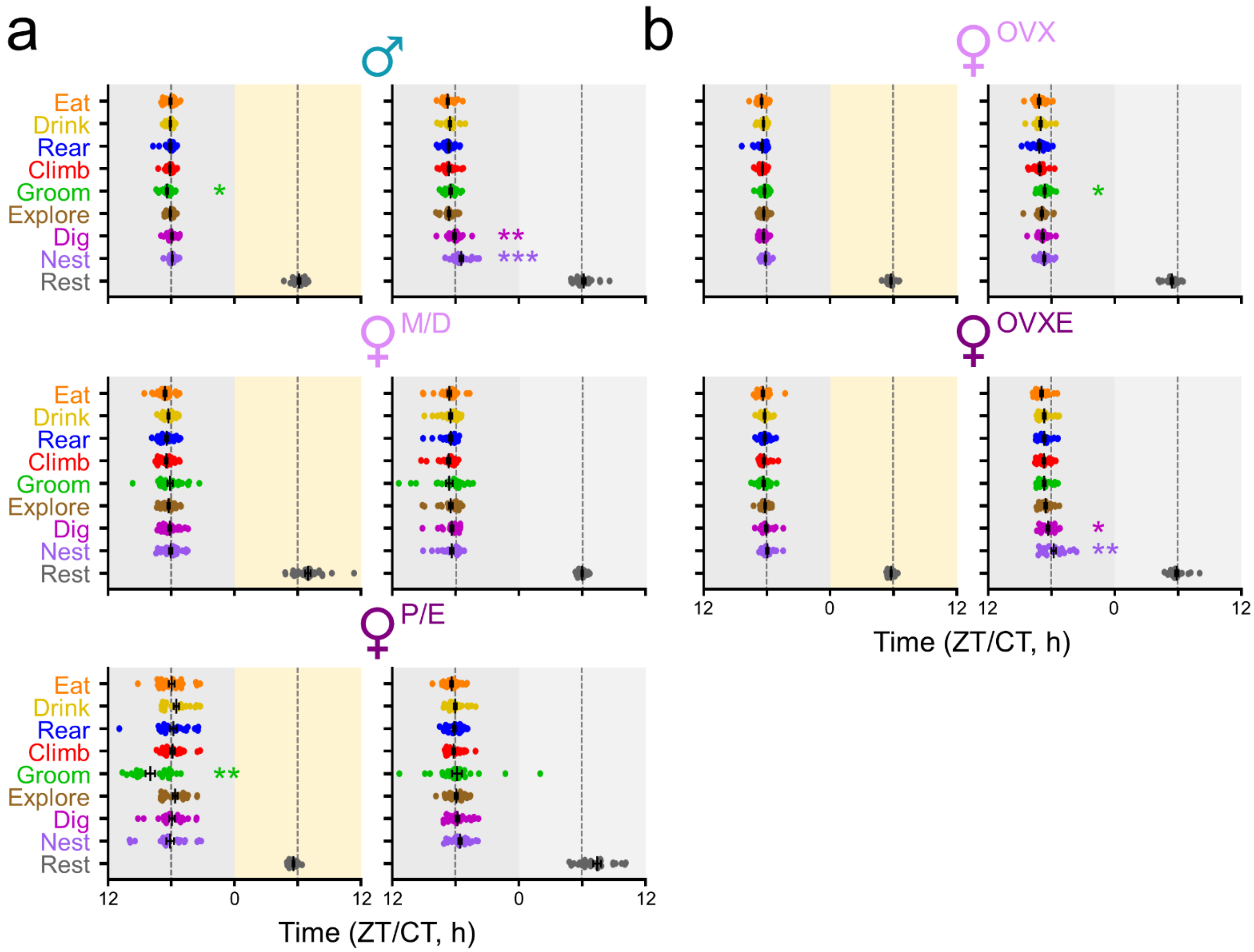


**Supplementary Figure 2. Sex and estrogen effects on behavior acrophases in individual mice. a,b)** Acrophases for **(a)** individual male, metestrus/diestrus (M/D) female, and proestrus/estrus (P/E) female mice, or **(b)** individual ovariectomized (OVX) and ovariectomized, estradiol-supplemented (OVXE) female mice, recorded in a 12h:12h light:dark (LD) cycle (left plots, gray and yellow shading) and in constant darkness (DD; right plots, gray and light gray shading). Each colored circle depicts an individual mouse's acrophase for a given behavior. Asterisks depict acrophases that differed significantly from those of other behaviors within groups. Lines and error bars depict mean ± SEM. One-way ANOVA with post-hoc Tukey's multiple comparisons test; *, $p < 0.05$; **, $p < 0.01$; ***, $p < 0.001$.

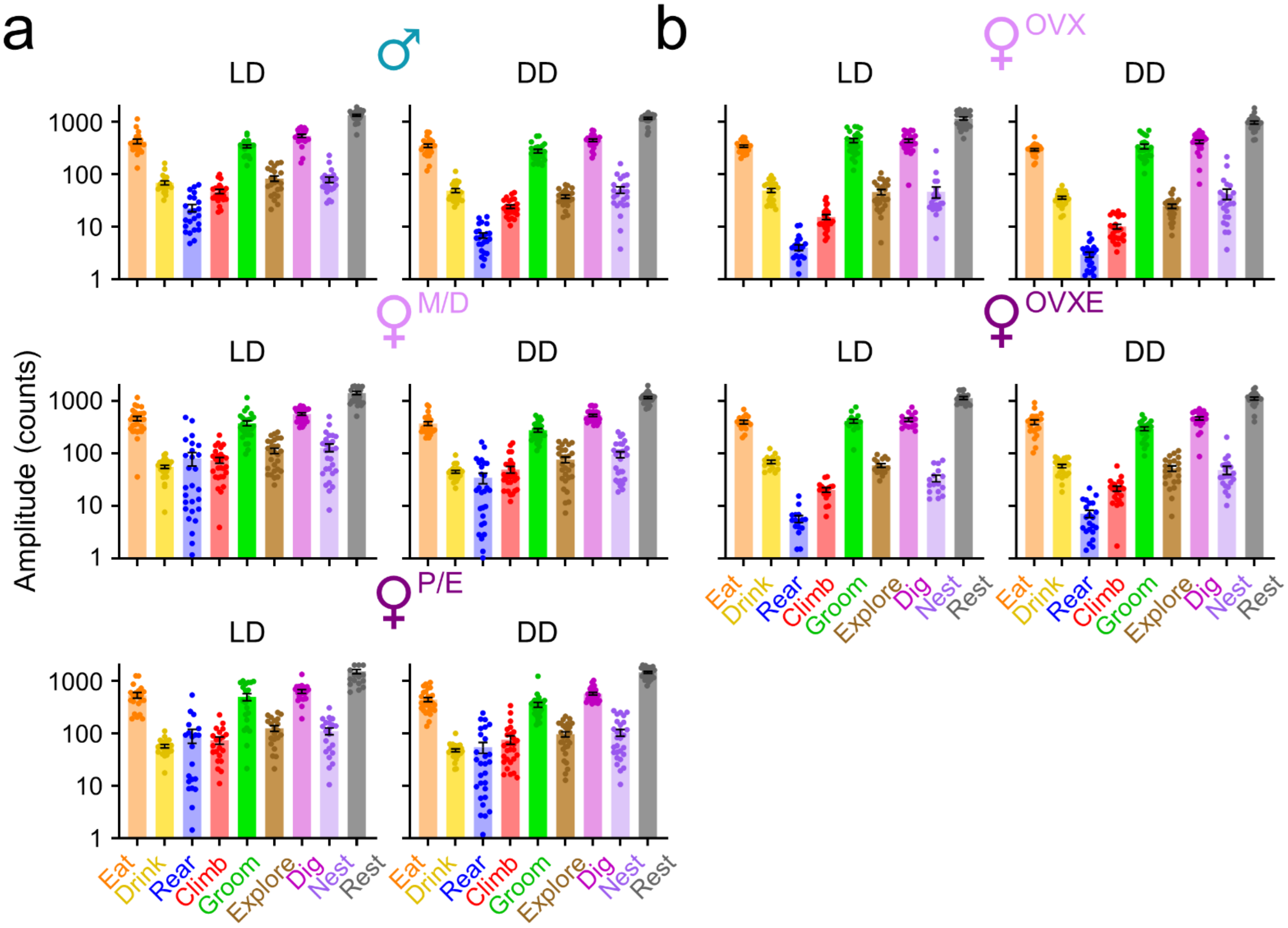


**Supplementary Figure 3. Sex and estrogen effects on absolute behavior amplitudes in individual mice.**

**a,b)** “Absolute” (non-normalized) amplitudes (in counts) for each behavior rhythm for **(a)** male, metestrus/diestrus (M/D), and proestrus/estrus (P/E) female mice, or **(b)** ovariectomized (OVX) and ovariectomized, estradiol-supplemented (OVXE) female mice, recorded in a 12h:12h light:dark (LD) cycle (left plots) and in constant darkness (DD; right plots).

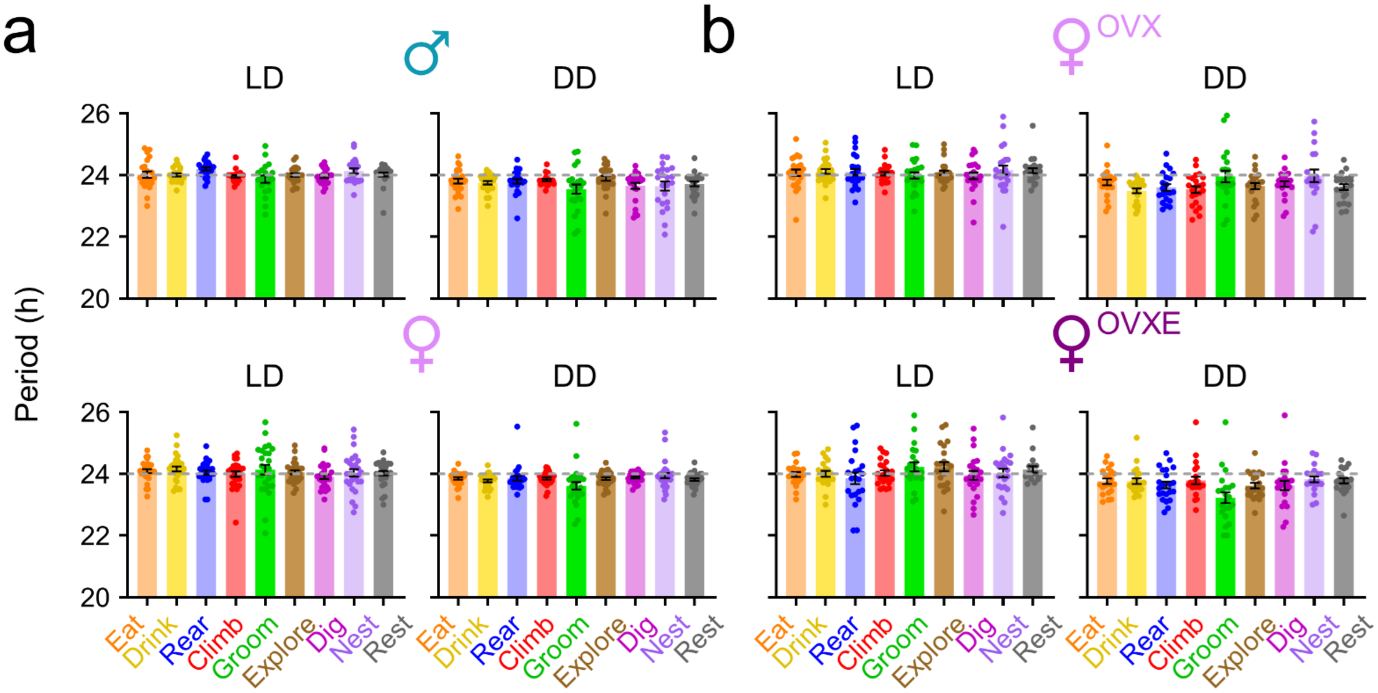


**Supplementary Figure 3. Sex and estrogen effects on period in individual mice. a,b)** Period for each behavior rhythm for **(a)** male and female mice, or **(b)** ovariectomized (OVX) and ovariectomized, estradiol-supplemented (OVXE) female mice, recorded in a 12h:12h light:dark (LD) cycle (left plots) and in constant darkness (DD; right plots).

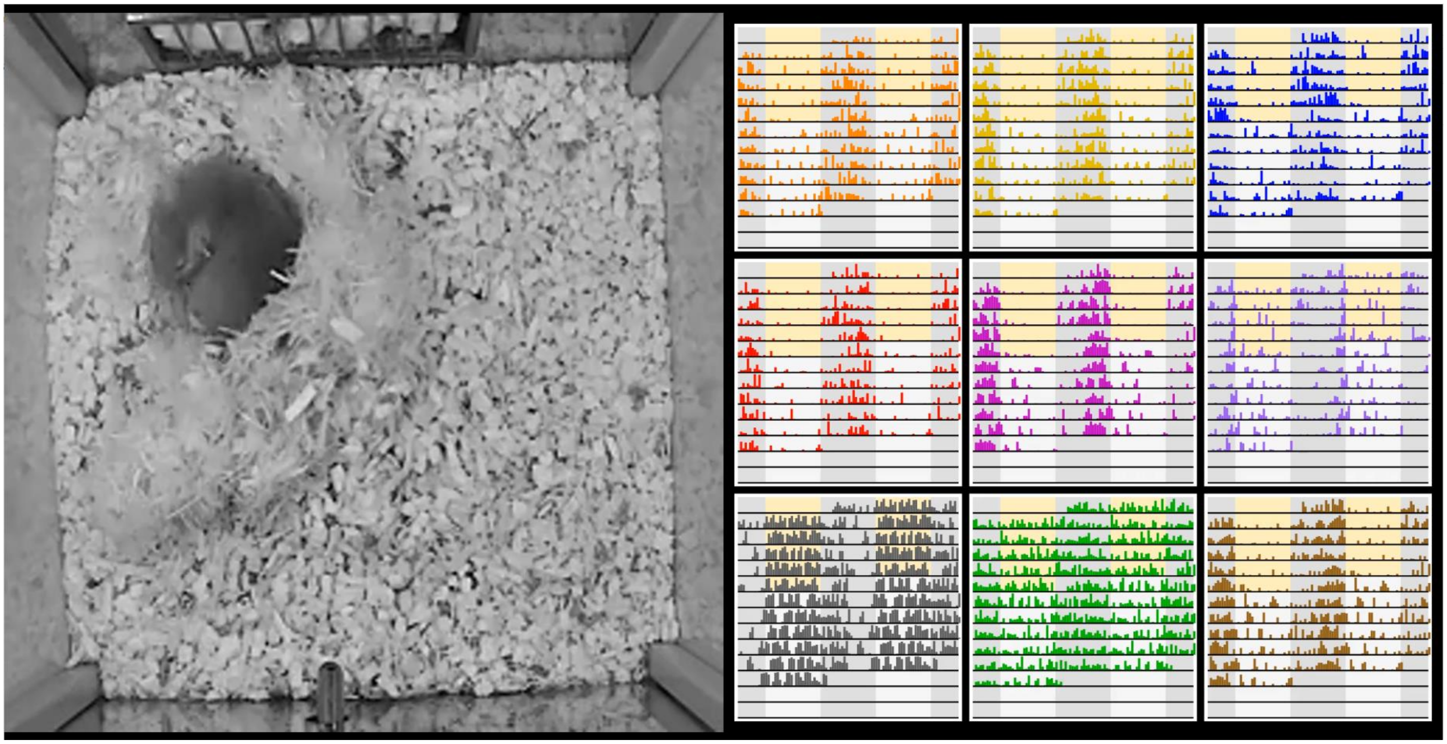

**Supplementary Video 1. Real-time video processing, behavior inference, and time series activity profile generation.**

| Behavior | Spatial Features | Temporal Features |
|---|---|---|
| Eat | Animal is positioned in front of the food hopper with its head in between food retention bars. | *Start* : Animal's head remains stationary or it moves forwards and backwards as it either remains within the food retention bars or shifts to be between neighboring food retention bars.<br>*Stop* : Animal moves head below the food hopper or makes a large movement across multiple food retention bars. |
| Drink | Animal's nose touches the end of the water spout. | *Start* : Animal's nose remains touching the end of the water spout or the animal is visibly licking the end of the spout.<br>*Stop* : Animal's nose detaches from the water spout. |
| Rear | Animal is not touching the side walls of the cage and its front paws are not touching the bottom of the cage. | *Start* : Animal remains stationary above the ground or hovers (side-to-side, forwards/backwards) without touching the ground or walls with its front paws.<br>*Stop* : Animal makes contact with the walls or base of the cage. |
| Climb | Animal is touching cage walls or cage objects to raise itself up from the bottom of the cage. | *Start* : Animal remains stationary on the wall or object, climbs higher up the wall or object, or jumps upward onto the wall or object.<br>*Stop* : Animal detaches from the wall or object or returns to the bottom of the cage. |
| Groom | Animal's head or paws are positioned such that they are interacting with its body. | *Start* : Animal licks, scratches, or investigates itself with occasional short pauses of $< 1$ s.<br>*Stop* : Animal's head and paws are not interacting with its body for a duration of $> 1$ s. |
| Explore | Animal is making broad movements across the cage. | *Start* : Animal moves across the cage without interacting with objects, bedding, or nesting material.<br>*Stop* : Animal interacts with any aspect of the cage (food hopper, water spout, walls, bedding, or nesting material). |
| Dig | Animal is positioned with its head downward into the cage bedding. | *Start* : Animal's head remains downward and stationary or actively moves the cage bedding.<br>*Stop* : Animal lifts its head from the bedding or begins to move nesting material instead of bedding. |
| Nest | Animal is touching nesting material. | *Start* : Animal actively moves nesting material with its head or paws.<br>*Stop* : Animal is no longer touching nesting material or moves nesting material without using its head or paws (i.e., on accident). |
| Rest | Animal is still with its head down; animal is in nest. | *Start* : Animal remains motionless for a period of $> 1$ s.<br>*Stop* : Animal moves head or body in any way. |

**Supplementary Table 1. Home cage behavior definitions.**
